# *De novo* draft assembly of the *Botrylloides leachii* genome provides further insight into tunicate evolution

**DOI:** 10.1101/152983

**Authors:** Simon Blanchoud, Kim Rutherford, Lisa Zondag, Neil J. Gemmell, Megan J. Wilson

## Abstract

Tunicates are marine invertebrates that compose the closest phylogenetic group to the vertebrates. This chordate subphylum contains a particularly diverse range of reproductive methods, regenerative abilities and life-history strategies. Consequently, tunicates provide an extraordinary perspective into the emergence and diversity of chordate traits. To gain further insights into the evolution of the tunicate phylum, we have sequenced the genome of the colonial Stolidobranchian *Botrylloides leachii*.

We have produced a high-quality (90 % BUSCO genes) 159 Mb assembly, containing 82 % of the predicted total 194 Mb genomic content. The *B. leachii* genome is much smaller than that of *Botryllus schlosseri* (725 Mb), but comparable to those of *Ciona robusta* and *Molgula oculata* (both 160 Mb). We performed an orthologous clustering between five tunicate genomes that highlights sets of genes specific to some species, including a large group unique to colonial ascidians with gene ontology terms including cell communication and immune response.

By analysing the structure and composition of the conserved gene clusters, we identified many examples of multiple cluster breaks and gene dispersion, suggesting that several lineage-specific genome rearrangements occurred during tunicate evolution. In addition, we investigate lineage-specific gene gain and loss within the Wnt, Notch and retinoic acid pathways. Such examples of genetic change within these highly evolutionary conserved pathways commonly associated with regeneration and development may underlie some of the diverse regenerative abilities observed in the tunicate subphylum. These results supports the widely held view that tunicate genomes are evolving particularly rapidly.

## Introduction

Tunicates are a group of marine suspension-feeding hermaphrodites found worldwide in the inter-or sub-tidal region of the seas. This subphylum of invertebrates is part of the Chordata phylum, phylogenetically positioned between the more basal Cephalochordata and the higher Vertebrata, of which they are considered the closest relatives (Fig. 1A; (Delsuc et al. 2006)). These organisms include a wide range of reproductive methods, regenerative abilities, developmental strategies and life cycles (Lemaire et al. 2008). Importantly, and despite a drastically different body plan during their adult life cycle, tunicates have a tissue complexity incipient to that of vertebrates (Fig. 1A), including a heart, a notochord, an endostyle and a vascular system (Millar 1971). In addition, this group of animals is undergoing rapid genomic evolution compared to higher vertebrates, with a greater nucleotide substitution rate observed in both their nuclear and mitochondrial genomes (Tsagkogeorga et al. 2010, 2012; Rubinstein et al. 2013; Berna and Alvarez-Valin 2014). Therefore, this chordate subphylum provides an excellent opportunity to study the origin of vertebrates, the emergence of clade specific traits and the function of conserved molecular mechanisms. Biological features that can be investigated in tunicates include, among others, the evolution of colonialism, neoteny, sessilness, and budding. However, there are currently only seven Tunicata genomes publicly available, of which three have been well annotated. There is thus a paucity in the sampling of this very diverse subphylum.

**Figure 1.**
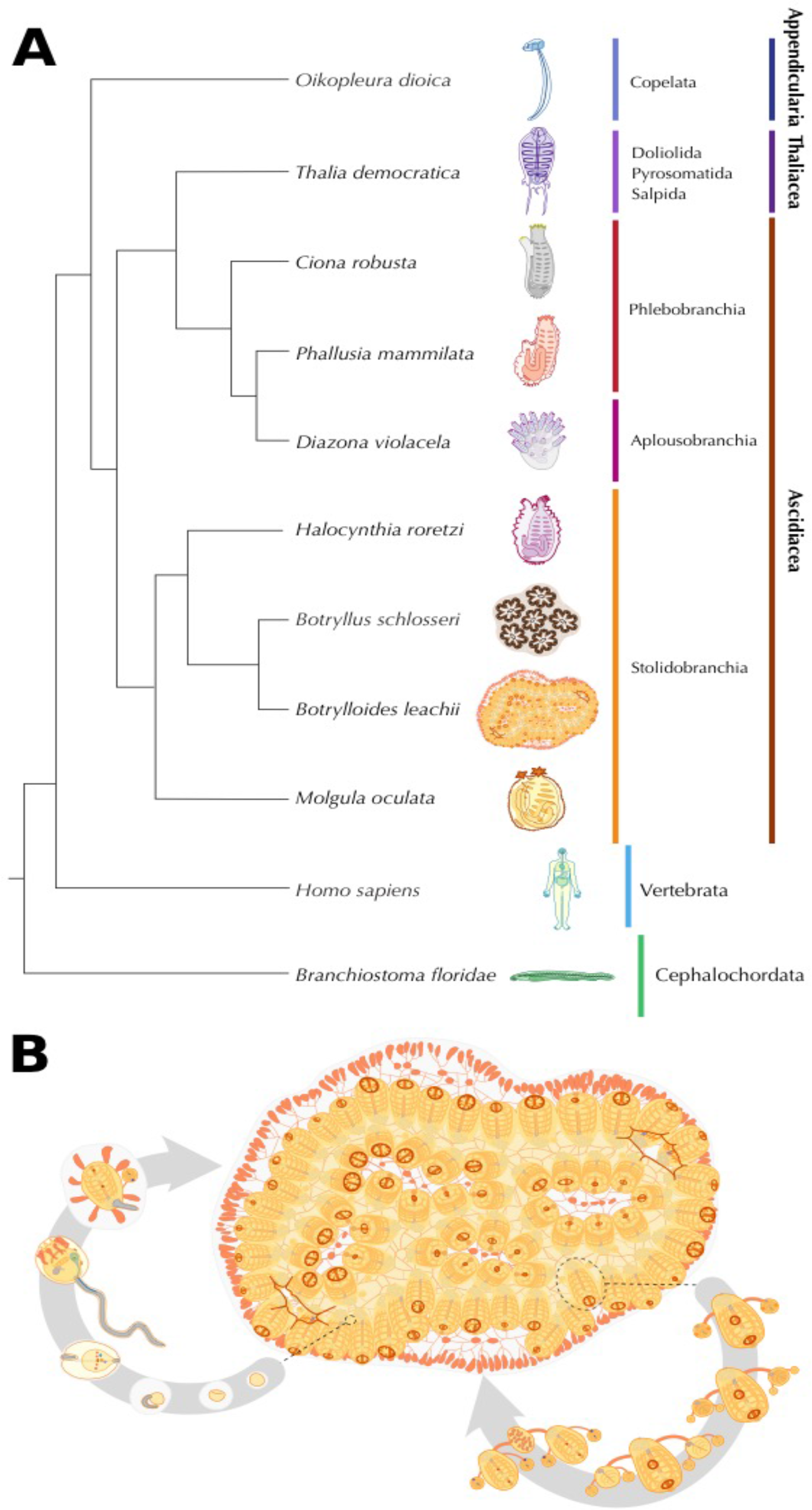
*B. leachii* phylogenetic position and life cycle. **A.** Schematic showing phylogeny of tunicates with respect to the chordate clade. **B.** Life cycle of *B. leachii*. The colony expands and grows by asexual reproduction (right loop). During favourable conditions such as warmer water temperatures, members of the colonies start sexual reproduction (left loop). The embryos remain with the colony in brood pouches until release. Hatched larvae attach to nearby substrates and begin metamorphosis into a zooid.

Tunicates are separated into seven orders contained in three classes (Fig. 1A): Appendicularia (order Copelata), Thaliacea (orders Pyrosomida, Salpida and Doliolida) and Ascidiacea (orders Aplousobranchia, Phlebobranchia and Stolidobranchia). Appendicularia is a class of planktonic free-swimming organisms that possess chordate traits common to all tunicate larvae including a notochord, neural tube and pharyngeal slits. These social neotenic animals form dioecious communities where each individual lives inside a special external mucous structure, termed house, which concentrates and funnels their food. *Oikopleura dioica* is the sole example of the Appendicularian to have its genome sequenced, showing exceptional compaction (70 Mb; (Seo et al. 2001)). Whether these animals can undergo regeneration has not yet been assessed.

Thaliacea is an order of planktonic pelagic animals forming cylindrical free-floating compound colonies (Piette and Lemaire 2015). These organisms can reproduce both sexually, through autogamy to initiate novel colonies, as well as asexually, through stolonial budding to increase the size of the colony. Owing to their peculiar life cycle and habitat, these tunicates have not been extensively studied, no genome has been sequenced and whether they can undergo regeneration remains unknown.

Ascidiacea consist of both solitary and colonial sessile benthic organisms. Solitary ascidians (Phlebobranchian and some families among the Stolidobranchian) reproduce sexually, releasing eggs through their atrial siphon for external fertilization, hence producing a motile larva. These larvae will colonize novel environments, attach to a submersed substrate and undergo metamorphosis into a sessile filter-feeding adult. These ascidians are capable of regenerating a limited set of organs, including their oral siphon (Auger et al. 2010) although regeneration capability reduces as they age (Jeffery 2015). These are currently the most sampled of the tunicates with five published genomes (*Ciona robusta* [formerly known as *C. intestinalis* type A; (Brunetti et al. 2015; Gissi et al. 2017)], *Ciona savigny, Molgula oculata*, *Molgula occulta*, *Molgula occidentalis;* (Dehal et al. 2002a; Small et al. 2007; Stolfi et al. 2014), two yet unpublished species (*Phallusia mamilata*, *Phallusia fumigata*; (Brozovic et al. 2016)) and two currently being assembled (*Halocynthia rorezi*, *Halocynthia aurantium*; (Brozovic et al. 2016)).

Colonial tunicates (Aplousobranchian and some families among the Stolidobranchian) are capable of both sexual, through autogamy, and asexual reproduction, through a wide range of budding types (palleal, vascular, stolonial, pyloric and strobilation; (Brown and Swalla 2012)). In addition, these compound organisms can undergo whole-body regeneration (WBR; reviewed in (Kürn et al. 2011)). Colonial ascidians are emerging as unique and increasingly popular model organisms for a variety of studies including immunobiology, allorecognition, angiogenesis and WBR (Rinkevich et al. 1995; Ballarin et al. 2001; Rinkevich et al. 2007a; Manni et al. 2007; Gasparini et al. 2008; Franchi et al. 2011; Lauzon et al. 2013; Rinkevich et al. 2013). However, colonial tunicates were only represented by *Botryllus schlosseri*, an ascidian that shows a significant expansion of its genome size when compared to the other available Tunicata genomes (725 Mb). To further investigate this fascinating subphylum and assess whether genome expansion is a prerequisite for coloniality and WBR, we have assembled and analysed the genome sequence of *Botrylloides leachii* (class Ascidiacea, order Stolidobranchia; (Savigny 1816)).

The viviparous colonial ascidian *B. leachii* can reproduce sexually through a tadpole stage that allows the settlement of a novel colony onto a new substrate (Fig. 1B). Each colony is composed of genetically identical adults (termed zooids) organized in ladder like systems and embedded in gelatinous matrix (tunic). While each adult has its own heart, they all share a common vascular system embedded within the tunic. In the presence of sufficient food supply, the size of the colony doubles approximately every 20 days through synchronized asexual reproduction, known as palleal budding. During this process each adult produces two daughter zooids that ultimately replace the mother, which is then resorbed by the colony. In addition, upon loss of all zooids from the colony, *B. leachii* can undergo whole-body regeneration and restore a single fully-functional adult in as little as 10 days from a small piece of its vascular system (Rinkevich et al. 1995). Furthermore, when facing unfavourable environmental conditions, these colonial tunicates can enter into hibernation, whereby all zooids undergo regression and are resorbed by the remaining vascular system. When a favourable environment is restored, mature adults will be restored to re-establish the colony (Burighel et al. 1976).

We have assembled and annotated the first *de novo* draft genome of the *B. leachii* by taking advantage of our recently published transcriptomes (Zondag et al. 2016). Using this genome, we have then undertaken a large-scale comparison of the four best-annotated ascidian genomes (*B. schlosseri*, *C. robusta, M. oculat*a and *O. dioica*) to gain insights into some of the diverse biological abilities that have evolved within the Tunicata.

## Results

### Genome assembly and annotation

To minimize contamination from marine algae and bacteria typically present in the pharyngeal basket of feeding *B. leachii*, we isolated genomic DNA from embryos of a single wild *B. leachii* colony. Genomic DNA was used to produce two libraries: one short-range consisting of 19,090,212 fragments (300 bp) of which 100 bp were paired-end sequenced - important for obtaining high coverage - and a second long-range mate pair with 31,780,788 fragments (1.5 - 15 kb size range, median ~3 kb) of which 250 bp were paired-end sequenced– important for scaffolding the assembly. Following quality checks, low quality reads were removed and sequencing adaptors were trimmed, thus resulting in a high-quality dataset of 86,644,308 paired-end and 12,112,004 single-end sequences (100 % with a mean Phred score > = 30, < 1 % with an adapter sequence, Fig. S1).

We then followed a reference-free genome characterization (Simpson 2014); provided with statistics from the human, fish (*Maylandia zebra*; (Bradnam et al. 2013)), bird (*Melopsittacus undulatus;* (Bradnam et al. 2013)) and oyster (*Crassostrea gigas*, (Zhang et al. 2012)) genomes for comparison; to estimate three properties of the *B. leachii* genome. First, k-mer count statistics were used to estimate the genome size to be 194.2 Mb (194,153,277 bp). This size is similar to that of the solitary *C. robusta, C. savigny and M. oculata* (160 Mb, 190 Mb and 160 Mb, respectively; (Dehal et al. 2002a; Small et al. 2007; Stolfi et al. 2014), larger than the compacted 70 Mb genome of *O. dioica* but appreciably smaller than the predicted 725 Mb genome of the related colonial ascidian *B. schlosseri*, of which 580 Mb have been sequenced (Voskoboynik et al. 2013a). Second, by quantifying the structure of the de Brujin graph obtained using the k-mer counts, the computational complexity of the assembly was estimated (sequencing errors 1/213, allelic differences 1/233, genomic repeats 1/2,439). With a cumulative occurrence of 1/106, the *B. leachii* genome is similar to that of bird, more variable than those of fish and human, but still quite less complex than the notably difficult oyster genome (Fig. S1). Third, sequence coverage was estimated using the distribution of 51-mers counts, showing a well-separated bimodal distribution with a true-genomic k-mers maximum at 31x coverage, similar to the human genome but higher than both the fish and the bird. Overall, these metrics suggest that *B. leachii* has a genome well suited for *de novo* assembly and that our sequencing could result in a high quality assembly.

*De novo* assembly using Metassembler (Wences and Schatz 2015) produced a genome of 159,132,706 bp (estimated percentage of genome assembled is 82 %), with an average sequencing coverage of 66x (after adaptor trimming). The assembly is composed of 1,778 scaffolds, with a N50 scaffold length of 209,776 and a L50 scaffold count of 223. The 7,783 contigs, with a N50 length of 48,085, and a L50 count of 781, represent a total size of 146,061,259 (92 %, Table 1). To evaluate the completeness of our assembly, we used the Benchmarking Universal Single-Copy Orthologs (BUSCO; (Simão et al. 2015)). This tool provides a quantitative measure of genome completeness by verifying the presence of a set of manually curated and highly conserved genes. Out of the 843 orthologs selected in metazoans, 760 (90 %) were found in our assembly of the *B. leachii* genome (File S1), a relatively high score when compared to the BUSCO score of frequently used genome assemblies such as *Homo sapiens* (89 %, GCA_000001405.15). In addition, we took advantage of our previous assembly of the *B. leachii* transcriptome (Zondag et al. 2016) to further assess the quality of our genome. Using BLAT (Kent 2002), we were able to map 93 % of transcript sequences (48,510/52,004) onto our assembly. Overall, these results indicate that our *de novo* genome is largely complete and suitable for annotation.

**Table 1.** *B. leachii* genome assembly statistics

|  |  |
| --- | --- |
| **Total length of assembly** | 159,132,706 bp |
| **Predicted genome size** | 194 Mb |
| **Number of scaffolds** | 1,778 |
| **Median scaffold length** | 43,485 bp |
| **N50 contig length** | 209,776 bp |
| **Estimated genome coverage before adaptor trimming** | 101x |
| **Estimated genome coverage after adaptor trimming** | 66x |
| **Number of predicted genes** | 15,839 |
| **% of the *B. leachii* reference transcriptome aligning to the genome** | 80 % |
| **% of *Ciona* proteins that have a significant match to the *B. leachii* genome** | 71% |
| **BUSCO score** | 90 % (760/843) |

*Ab initio* genome annotation was performed using MAKER2 (Holt and Yandell 2011) and predicted 15,839 coding genes, of which 13’507 could be classified using InterProScan (Jones et al. 2014). Comparing these predictions with our mapping of the transcriptome, we found out that 83 % of our aligned cDNA (40,188/48,510) mapped to a predicted gene locus thus spanning 78 % of the annotated genes (12,395/15,839). In addition, a total of 4,213 non-coding sequences were predicted using Infernal (Nawrocki and Eddy 2013), Snoscan (Lowe 1999) and tRNAscan-SE (Lowe 1999). Finally, repetitive elements were annotated using RepeatMasker (Smit et al. 2015) and a species-specific library created using the RepeatModeler module (Smit and Hubley 2015). Eighteen percent of the genome was identified as containing repetitive elements (Table 2 and Table S1), a majority (17%) of these being interspersed repeats. This proportion is similar to that found in other tunicates including *C. robusta* (17%), M. oculata (22%) and *O. dioica* (15%), while being lower than that in *B. schlosseri* (60%).

**Table 2.** Comparison of sequenced tunicate genomes and their most prominent biological features.

|  | *Ciona robusta* | *Botrylloides leachii* | *Botryllus schlosseri* | *Oikopleura dioica* | *Molgula oculata* |
| --- | --- | --- | --- | --- | --- |
| **Genome size** | 160 Mb | 194 Mb | 725 Mb | 72 Mb | 160 Mb |
| **Number of sequences** | 4,390 | 1,778 | 121,094 | 1,260 | 10,554 |
| **Fraction of repetitive DNA** | 17% | 18% | 60% | 15% | 22% |
| **Predicted gene number** | 16,671 | 15,839 | 27,463 | 16,749 | 15,313 |
| **GC content** | 36% | 41% | 41% | 40% | 36% |
| **Body structure** | solitary, sessile | colony, sessile | colony, sessile | solitary, motile | solitary, sessile |
| **Reproduction** | sexual, hermaphrodite | asexual sexual, hermaphrodite | asexual sexual hermaphrodite | sexual, separate sexes | sexual, hermaphrodite |
| **Regenerative ability** | specific organs | WBR | WBR | unknown | unknown |

To further characterize the genome of *B. leachii*, we set out to compare it to four available Tunicata genomes. First, we quantified the number of sequences from their proteomes which mapped onto our assembly using tBLASTn (Camacho et al. 2009): *C. robusta* 71% (10,507/14,740), *M. oculata* 77 % (12,788 / 16,616), *B. schlosseri* 71 % (32,944/ 46,519) but only 30% for *O. dioica* (9,009/29,572). Secondly, we performed an all-to-all search for protein orthologs using the OrthoMCL clustering approach (Li et al. 2003) to identify any orthologs between the tunicate genomes (Fig. 2A). Clustering the combined protein set from all five genomes resulted in 17,710 orthologous groups. By classifying each group based on which tunicate has proteins in it, we identified the presence of five larger set of orthologs: those shared by all species (17% of all groups), those shared by all sessile tunicates (11%), those between the two colonial species (11%) and two groups unique to *B. schlosseri* and *O. dioica* (15 % and 12 %, respectively; Fig. 2A). Thirdly, the proteins specific to an organism were removed from the corresponding proteome, and a new mapping to the *B. leachii* genome was performed. Mapping of the proteome using only the conserved sequences is 93 % for *B. schlosseri* and 45 % for *O. dioica*.

**Figure 2.**
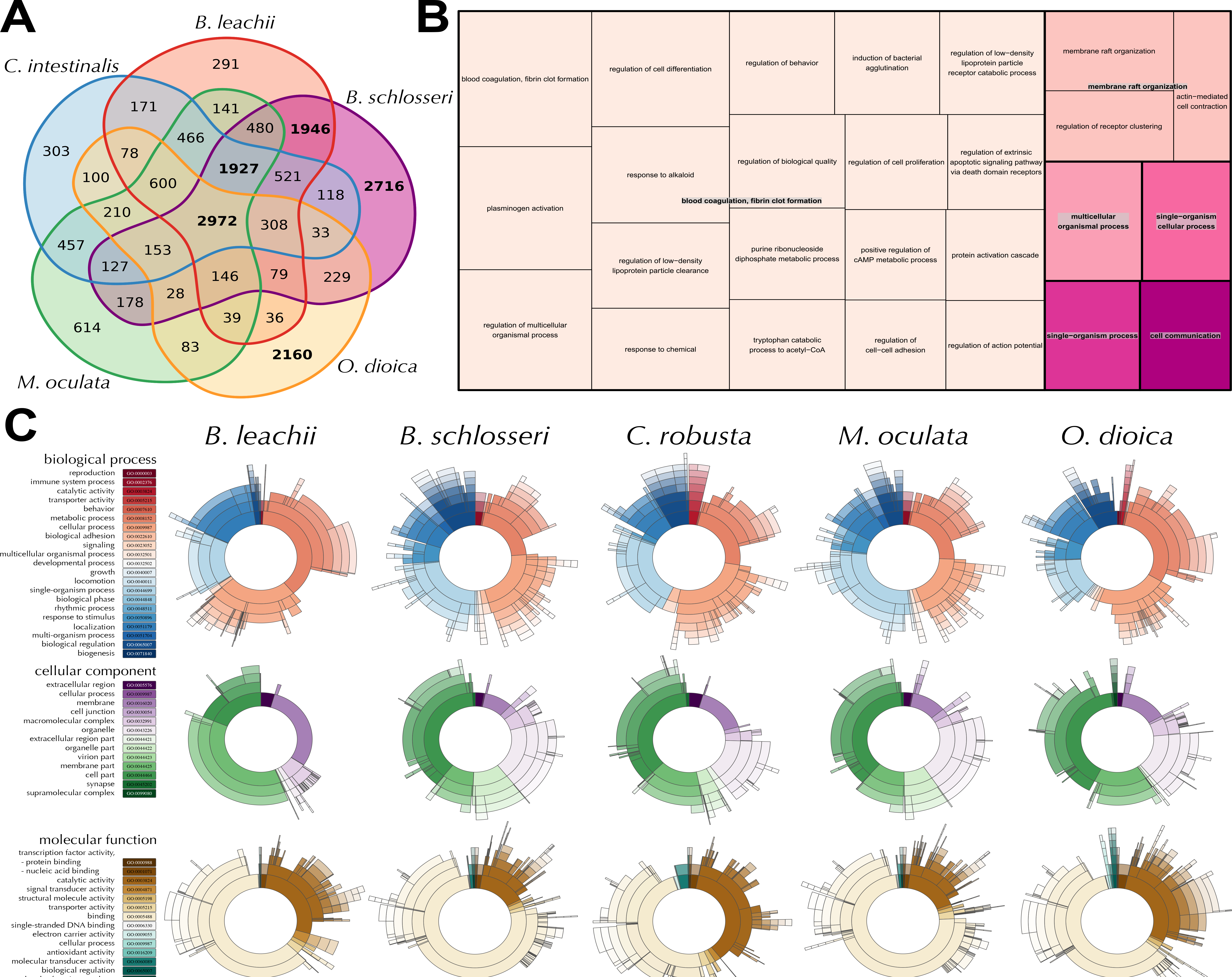
(Following page). Comparison of the tunicate genomes. **A.** Clustering of orthologous protein sequences. Indicated are the number of cluster groups, each of which contains at lest two proteins. **B.** Treemap representation of the overrepresented GO Biological Processes terms within the ortholog groups shared between *B. leachii* and *B. schlosseri* genomes but not with *C. robusta, O. dioica* and *M. oculata*. Each rectangle size is proportional to the p-value for the GO term. **C.** Distribution of the three classes of GO terms for each species. The color-codes (left) are common for the entire row.

To get insights into the potential biological function underlying these ortholog groups, we analysed the distribution of Gene Ontology (GO) terms for each cluster (Fig. S2). Interestingly, an important fraction of the shared orthologs are related to G-protein signalling, a conserved family of proteins involved in a variety of transduction mechanisms (Iwasa et al. 2003; Philips et al. 2003; Murata et al. 2001). Additionally, given that these proteins are potentially novel to colonial tunicates, a cross-species approach for GO enrichment was performed using the Human GO database as background (Fig. 2B, (Primmer et al. 2013)). Finally, we compared the composition of GO terms at the genomic level (Fig. 2C). Despite *B. schlosseri* having a larger predicted gene number compared to the other analyzed tunicates, the overall proportion of GO groups terms were distributed similar between all genomes (Fig. 2C), indicating no expansion of one particular functional group in *B. schlosseri*. Overall, these analyses showed that our assembly and annotation is consistent with the other tunicate genomes and provides novel insights into their shared mechanisms and potentially for evolutionarily conserved mechanisms as well.

### Ancient gene linkages are fragmented in tunicate genomes

Ancient gene linkages are highly conserved sets of genes that are spatially restricted, commonly occurring in clusters (Garcia-Fernàndez 2005). These clusters arose in a common ancestor and were preserved because of a common regulatory mechanism such as cis-regulatory elements located within the cluster. The homeobox-containing *Hox* gene family, typically composed of 13 members in vertebrates (Hoegg and Meyer 2005), is among the best-studied examples of such an ancient gene cluster and is critical for the correct embryonic development (Pearson et al. 2005). The linear genomic arrangement of genes within the *Hox* cluster reflects their spatial expression along the anterior-posterior body axis (Pascual-Anaya et al. 2013), which establishes regional identity across this axis. The basal cephalochordate *B. floridae* has all 13 *hox* genes located in a single stereotypical cluster, along with an additional 14^th^ gene (Fig. 3B; (Takatori et al. 2008)), suggesting that the chordate ancestor also had an intact cluster. However in tunicates, this clustering appears to be lost. In *C. robusta*, the nine identified *Hox* genes are distributed across five scaffolds, with linkages preserved only between *Hox2, Hox3* and *Hox4; Hox5* and *Hox6*; *Hox12* and *Hox13* (Fig. 3; (Spagnuolo et al. 2003; Wada et al. 2003)). In *O. dioica*, the total number of *Hox* genes is further reduced to eight, split between 6 scaffolds, including a duplication of *Hox9* (Fig. 3A; (Edvardsen et al. 2005)). In *M. oculata* we could identify only six *Hox* genes, divided between 4 scaffolds, with clustering retained for the *Hox10*, *Hox11* and *Hox12* genes (Fig. 3). In Botryllidae genomes, the same seven *Hox* genes are conserved (Fig 3B), with a preserved linkage between *Hox10*, *Hox12* and *Hox13* in *B. leachii* and three copies of *Hox5* present in *B. schlosseri*. Altogether, the separation of the tunicate *Hox* cluster genes supports the hypothesis that reduction and separation of this ancient gene linkage occurred at the base of tunicate lineage (Edvardsen et al. 2005). In addition, *Hox9* appears to be specifically retained in neotenic Tunicates while there is no pattern of conserved *Hox* cluster genes specific to colonial ascidians.

**Figure 3.**
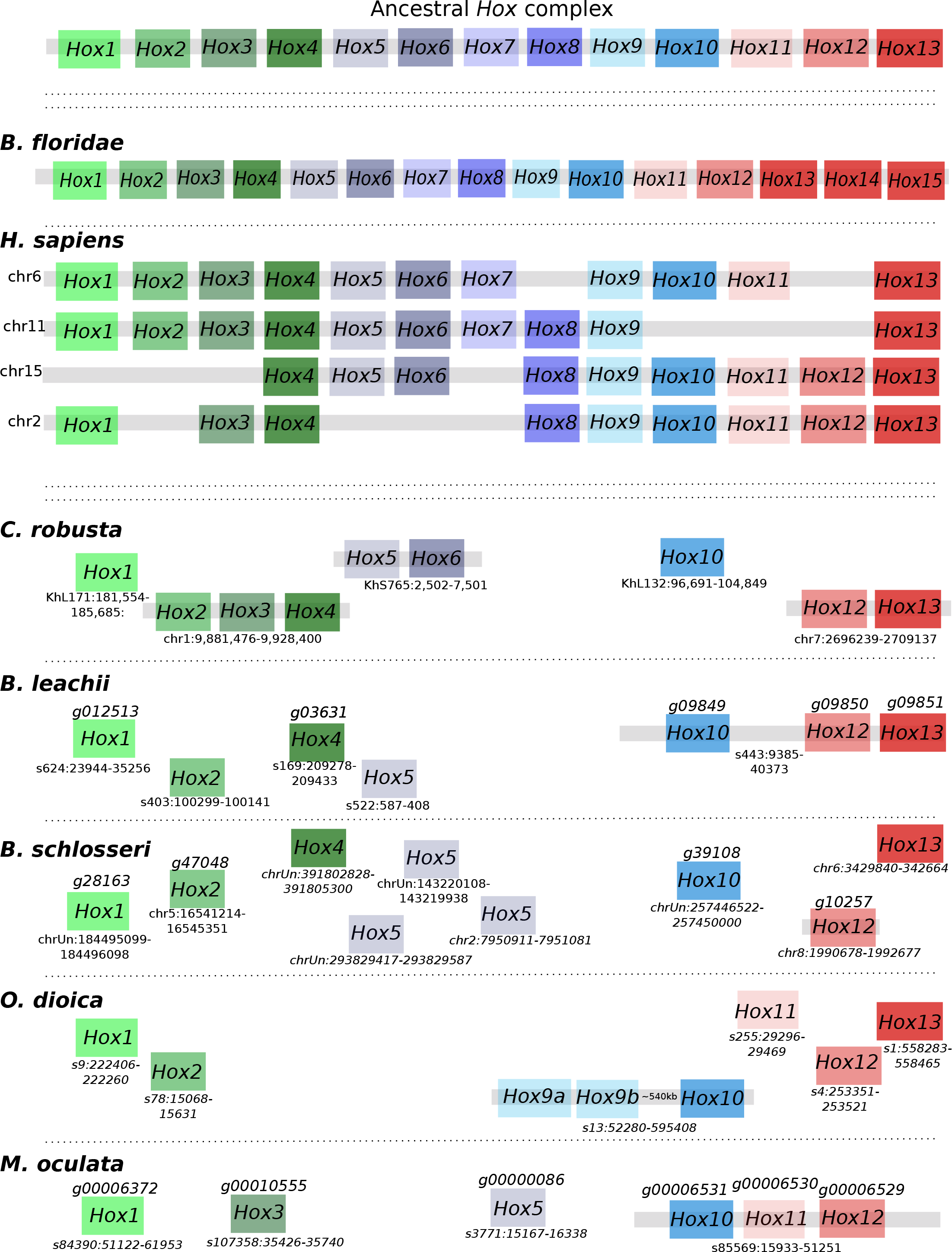
*Hox* genes are dispersed and reduced in number within tunicate genomes. Schematic depicting *Hox* gene linkages retained in five tunicate genomes in comparison to the ancestral *Hox* complex, which included thirteen genes. Orthologous genes are indicated by common colours. Chromosome or scaffold number is shown, along with gene ID when available for newly annotated genomes.

A second ancient homeobox-containing gene linkage is the *NK* cluster. This cluster, predicted to be present in the last common ancestor of bilaterians (Luke et al. 2003), consists of *Msx, Lbx*, *Tlx*, *NKx1, NKx3, NKx4* and *NKx5* (Fig. 4). In *B. floridae*, linkages between *Msx, NKx4* and *NKx3*; as well as between *Lbx* and *Tlx* provide evidence of retained ancestral clustering while *NKx5* was lost (Fig. 4; (Luke et al. 2003)). However in vertebrates, NKx5 is still present, while only the gene linkages between *Lbx* and *Tlx* as well as between *Nkx4* and *Nkx3* remain (Fig. 4; (Garcia-Fernàndez 2005)). To further clarify the evolution of this ancestral cluster in tunicates, we determined the structure of the NK cluster within five ascidian genomes. In all these species, *NKx1* is absent and no evidence of clustering could be found with all identified orthologs located on different scaffolds (Fig. 4). In *C. robusta*, *M. oculata* and *O. dioica* only five members of this cluster remain, with the loss of either *Lbx* or *Tlx* as well as of *NKx3* and the duplication of the ortholog of *NKx4* (Fig. 4). By contrast, in the colonial tunicates *B. leachii* and *B. schlosseri*, *Tbx*, *Lbx* and *NKx3* are all present. In *B. schlosseri*, *Msx1* is absent and *NKx4* duplicated. In the *B. leachii* genome, *NK1* is the only ancestral cluster member to be missing and *Nk5* has been duplicated (Fig. 4). Altogether, these results suggest that there has been a loss of *NKx5* in Cephalochordate, one of *NKx1* in Tunicate and that the combination of *NKx3*, *Lbx* and *Tbx* may be specific to colonial ascidians.

**Figure 4.**
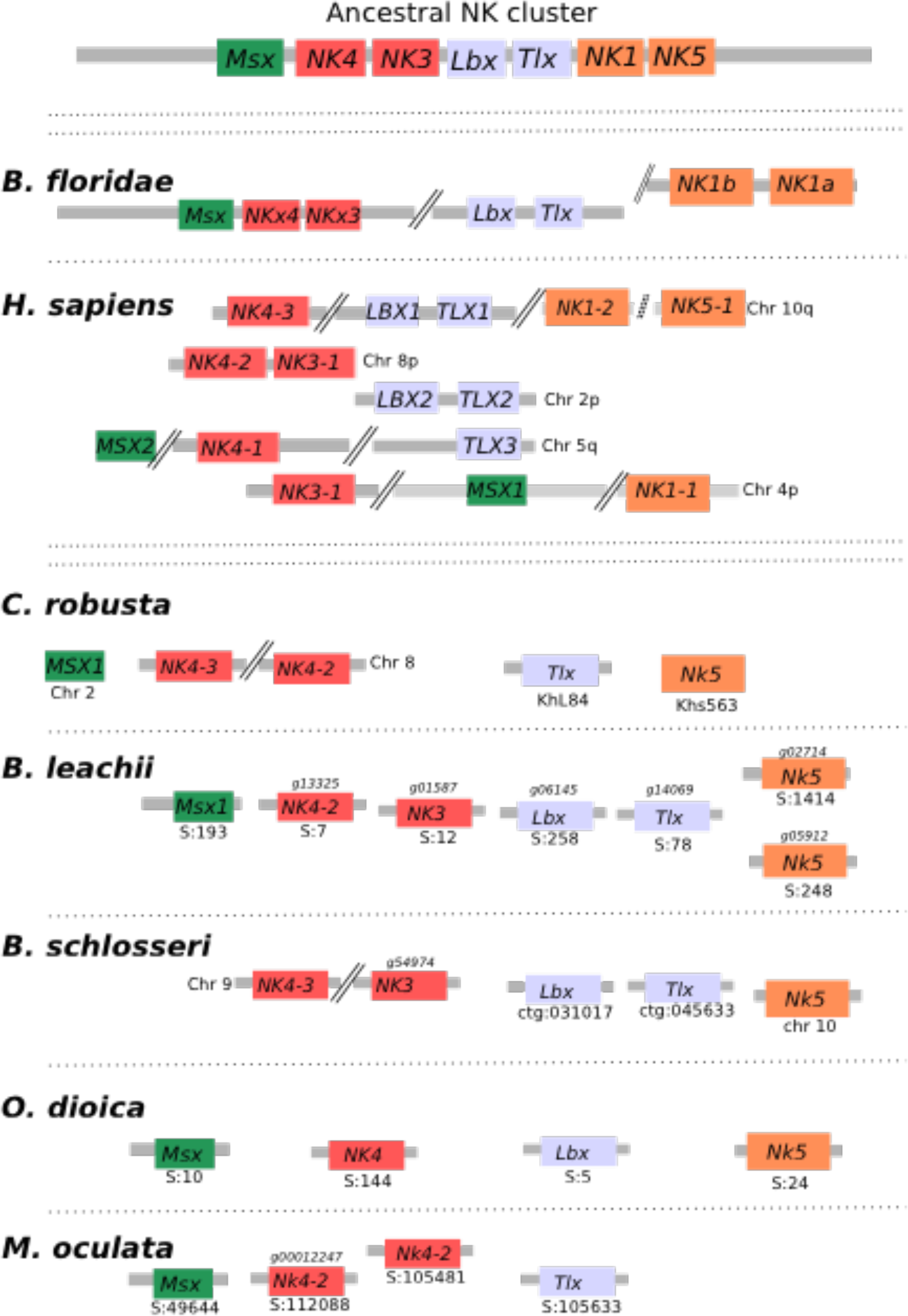
NK homeobox cluster genes are fragmented within tunicate genomes. Schematic depicting the organization of the *NK homeobox* cluster genes among the studied chordate genomes. Double-parallel lines indicate > 1Mb distance between genes. Chromosome or scaffold number is shown, along with gene ID when available for newly annotated genomes. Orthologous genes are indicated by common colours.

A third ancient linkage that we investigated is the pharyngeal cluster, a gene group present in hemichordates, echinoderm and vertebrates genomes that is considered to be Deuterosome specific (Simakov et al. 2015). The cluster groups *foxhead domain protein* (*FoxA*), *NKx2 (NKx2.2 and Nkx2.1)*, *Pax1/9*, *mitochondrial solute carrier family 25 member 21* (*slc25A21)*, *mirror-image polydactyly 1 protein* (*mipol1*), *egl nine homolog 3* (*egln3*) and *dehydrogenase/reductase member* 7 (*dhrs7*). Among these, s*lc25a21, Pax1/9*, *mipol1* and *FoxA* pairs are also found in protostomes suggesting an even more ancient origin (Simakov et al. 2015). The pharyngeal cluster is thought to have arisen due to the location of the regulatory elements of *Pax1/9* and *FoxA* within the introns of *slc25A21* and *mipol1* (Santagati et al. 2003; Wang et al. 2007), constraining the genes to remain in tight association with each other. In the *B. floridae* genome, the entire cluster is located on the same scaffold, with the exception of the *Nkx2.1* and *Nk2.2* gene pair located on separate scaffold. In *C. robusta*, only orthologs of *FoxA*, slc25a29, *Pax1* and *Pax9* could be identified. Nevertheless, all of them are located on the same chromosome (Fig. 5). In *O. dioica*, the cluster appears even further reduced. While orthologs of *FoxA*, *Pax1/9* and *Nkx2.2* genes were found on different scaffolds, only one rather distant linkage (> 1 Mb) between *a Pax-like* gene and *slc25A21* is retained. For both *B. schlosseri* and *M. oculata*, there was no evidence of clustering between genes (Fig. 5). In the *B. leachii* genome, *mipol1* is the sole missing gene from this cluster. However, only the pairing of a *Pax-like* and *slc25A21* genes remains (Fig. 5). Altogether, these results suggest that most of the Tunicates did not conserve the structure of this ancient linkage, but it is unknown what consequences this would have to their expression and function.

**Figure 5.**
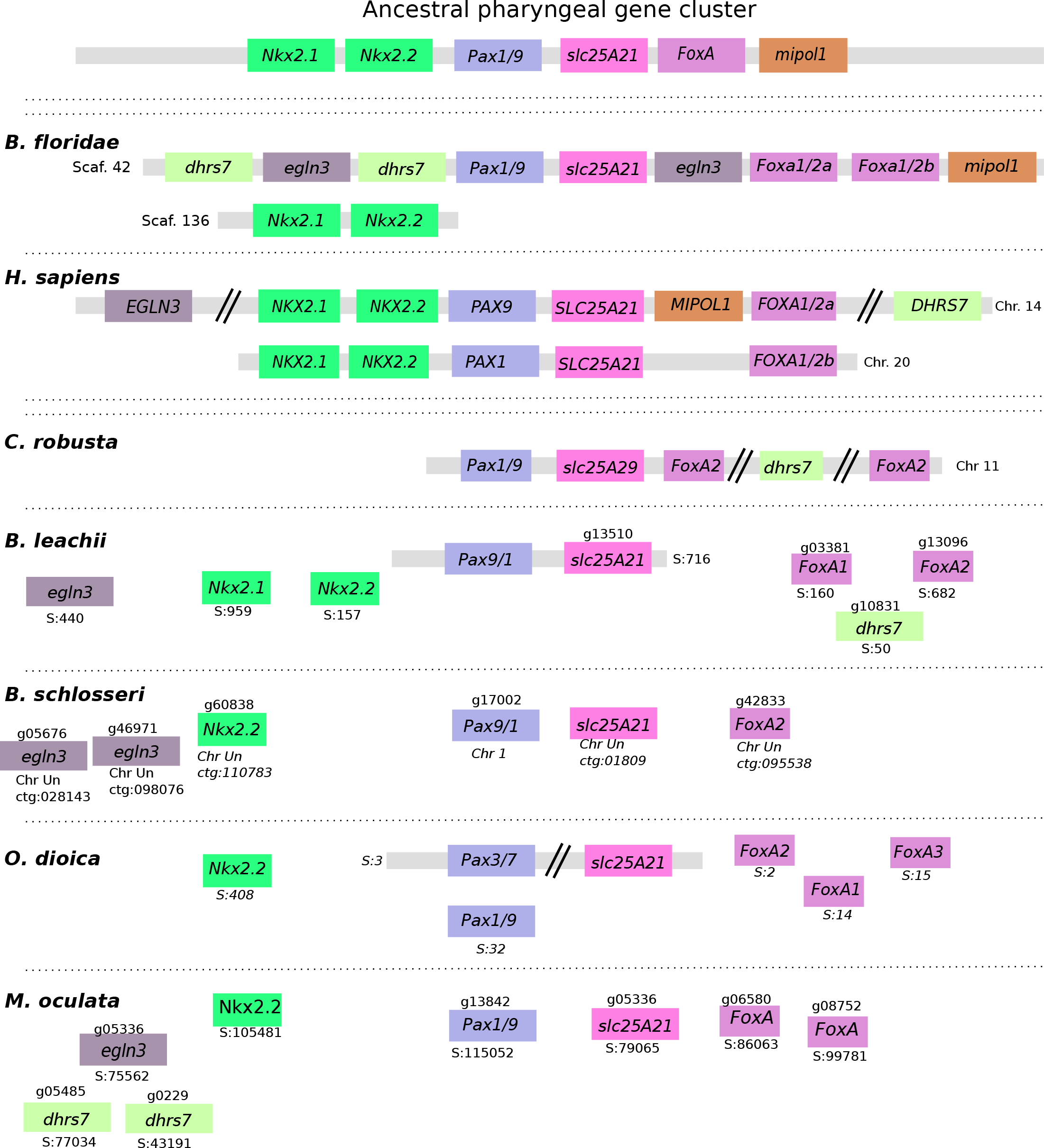
Ancestral gene linkages remain between a few pharyngeal cluster genes in tunicate genomes. Gene order of the six pharyngeal cluster genes, *NK2.1, NK2.2, Pax1/9* and *FoxA* in chordate genomes. Double-parallel lines indicate > 1 Mb distance between genes. Chromosome or scaffold number is shown, along with gene ID when available for newly annotated genomes. Orthologous genes are indicated by common colours.

### Lineage-specific changes to cell-signalling pathways in Botryllidae genomes

To dissect more specifically the evolution of colonial ascidians, we examined the genomes of *B. leachii* and *B. schlosseri*, looking for key components of signalling pathways required for metazoan development and regeneration. Of particular interest, we focused on the Wingless-related integration site (Wnt), Notch and Retinoic acid (RA) signalling pathways. All three of these pathways have been implicated in WBR and asexual reproduction in colonial tunicates (Rinkevich et al. 2008, 2007b; Zondag et al. 2016).

### Wnt pathway

Wnt ligands are secreted glycoproteins that have roles in axis patterning, morphogenesis and cell specification (Loh et al. 2016). The ancestral gene family appears to originate very early on during multi-cellular evolution and to be composed of eleven members (Kusserow et al. 2005; Guder et al. 2006). The *Wnt* gene family expanded to 19 members in the human genome, while independent gene loss has reduced this family to 7 genes in *Drosophila melanogaster* and *Caenorhabditis elegans* (Prud’homme et al. 2002). Consequently, we set out to investigate whether the *Wnt* gene family has either expanded or contracted during Tunicata speciation.

We found an increase in the number of *Wnt5a* genes among Styelidae genomes. In *B. schlosseri*, we identified 15 *Wnt* members, including seven *Wnt5a-like* genes on multiple scaffolds (Fig. 6, Table S2). In the *B. leachii* genome, fourteen *Wnt* ligand genes were identified, including four *Wnt5a* genes located on the same scaffold near *Wnt4* (Fig. 6). *M. oculata* has only 7 Wnt ligand genes, including three *Wnt5a-like* genes (Fig. 6, Table S2). In comparison, *C. robusta* has a total of 11 *Wnt* genes, including a single *Wnt5a* gene (Fig. 6, Table S2; (Wada et al. 2003)). In the compact *O. dioica* genome, this number has reduced to 6 (Wnts 3, 4, 7, 11 and 16), none of which are *Wnt5a* orthologs (Table S2). Overall, this suggests that an expansion through gene duplication of the Wnt5 family occurred during tunicate evolution, but was lost in some lineages.

**Figure 6.**
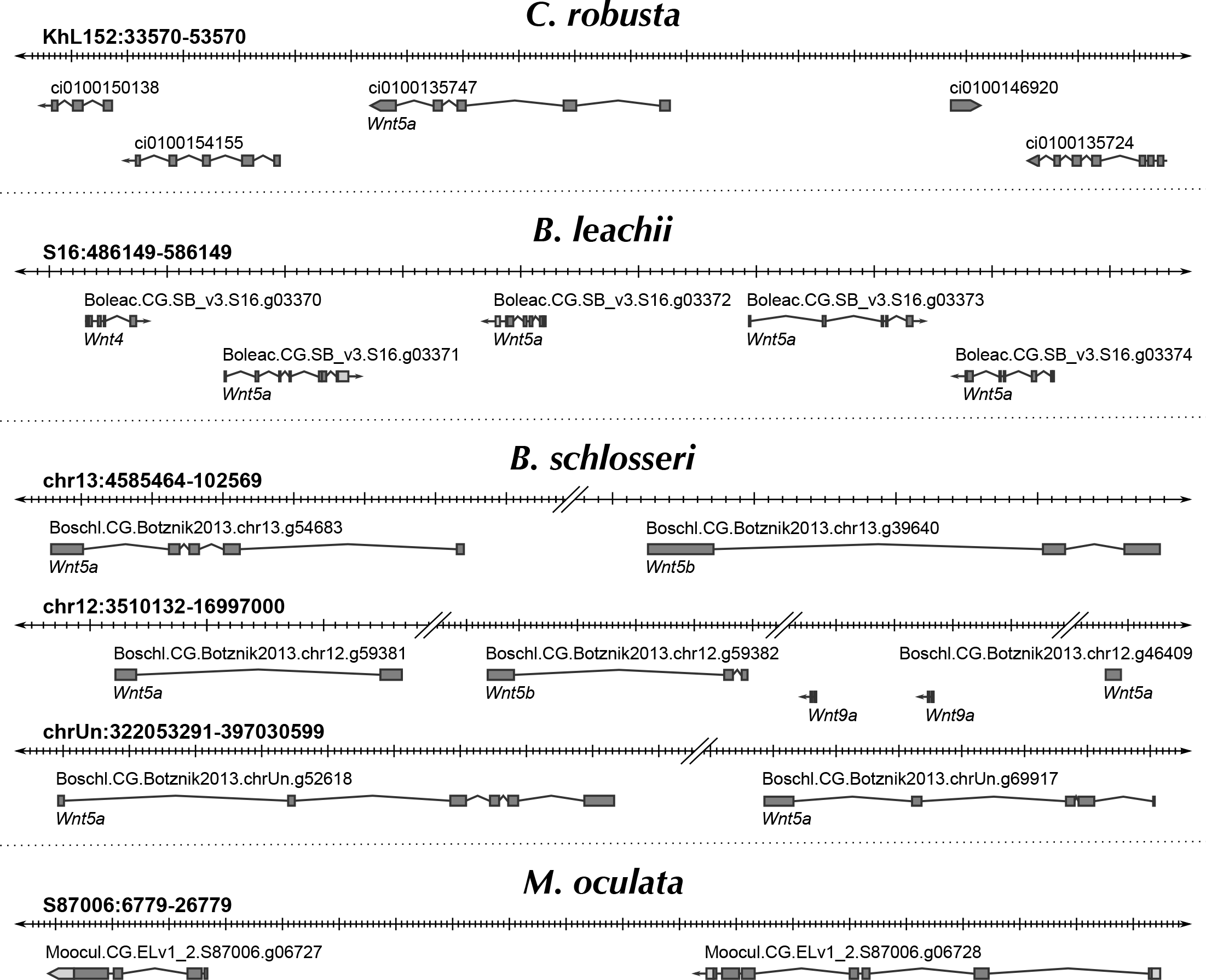
Duplication of *Wnt5a* genes in tunicate genomes. Schematic showing the genomic location of *Wnt5*-like genes within each indicated genome. Note that no *Wnt5a* ortholog is present in the *O. dioica* genome. Double-parallel lines indicate> 1Mb distance between genes.

To assess the functionality of the Wnt pathway in Tunicates, we set out to assess whether its downstream effectors are themselves present in the available genomic data. The downstream pathways activated by Wnt ligands are divided into canonical, non-canonical calcium and non-canonical planar cell polarity. The Wnt5a ligand is associated with both of the non-canonical pathways through binding of membrane receptors that include *frizzled* (*Fzd4*), *receptor tyrosine kinase-like orphan receptor 1/2* (*Ror1/2*) and *atypical tyrosine kinase receptor* (*Ryk*). Further downstream, disheveled (dsh), β-catenin (Cnntb), Axin, low-density lipoprotein receptor-related protein 5/6 (LRP5/6) and nuclear factor of activated T-cells (NFAT) are proteins essential for triggering intracellular responses to Wnt signalling (MacDonald et al. 2009). We identified orthologs for each of these signalling transduction molecules in all Tunicata genomes (Table S2), with no evidence of further gene duplication events. This supports the interpretation that signalling through the Wnt pathway is functional in tunicates.

### Notch pathway

Notch receptors are transmembrane proteins that are involved in cell-cell signalling during development, morphogenesis and regeneration (Hamada et al. 2015). Following activation through the binding of the delta or jagged/serrate ligands, the intracellular domain of Notch is cleaved and induces the expression of downstream target genes including the *hes (hairy and enhancer of split*) gene family members (Guruharsha et al. 2012). The presence of both Notch and the Delta/Serrate/lag-2 (DSL) proteins in most metazoan genomes suggests that their last common ancestor had a single copy of each gene (Gazave et al. 2009). To establish how this pathway has evolved in tunicates, we screened these genomes for the Notch receptor using the conserved lin-Notch repeat (LNR) domain, and for genes encoding probable Notch ligands such as genes from the DSL family.

In all examined genomes, only a single *Notch* receptor gene was identified while the number of ligand genes varied (Table S3). The *C. robusta* genome contains two *DSL* genes, while *O. dioica, M. oculata* and *B. schlosseri* possess only a single *DSL*. By contrast, we found three DSL genes in *B. leachii* (Table S3). To determine the relationships between these identified tunicate DSL-like genes, a phylogeny was constructed along with other chordate DSL proteins. All three *B. leachii* genes are Delta orthologs, two of them related to the *B. schlosseri* and *Cionidae* copy; the third one closer to the *M. oculata* and *H. roretzi* variant. The mouse, human and zebrafish delta and delta-like (DLL) proteins form a discrete clade loosely related to the genes found in Cephalochordates and Tunicates (Fig. 7, shaded box). Jagged proteins form a separate clade (Fig. 7). The tunicate DSL-like proteins present long phylogenetic branches, suggestive of greater diversity, which is also observed in the protein alignment (Fig. S3). This suggests that the tunicate DSL proteins are diverging rapidly from each other, indicative of lineage specific evolution of DSL-like genes.

**Figure 7.**
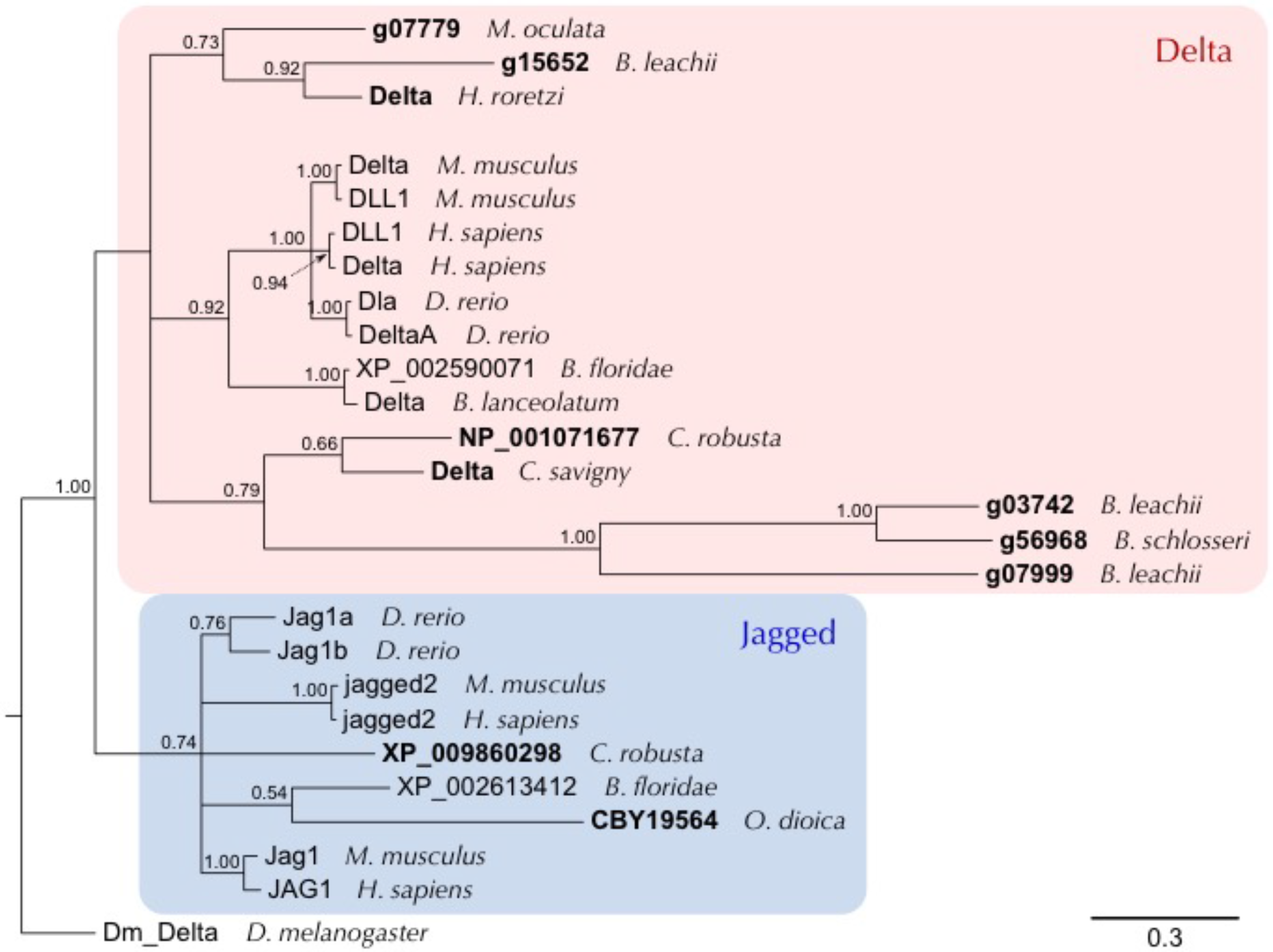
*B. leachii* Notch pathway. Bayesian phylogenetic tree depicting the relationship between tunicate and vertebrate DSL proteins, using *Drosophila* Delta to root the tree. Tunicate proteins are shown in bold and shaded areas correspond to Delta and Jagged groupings. Branch support values (probabilities) are indicated.

### Retinoic acid signalling

Retinoic acid (RA) is an extracellular metabolite that is essential for chordate embryonic development. RA is synthesized from retinol (vitamin A) by two successive oxidation steps. In the first step, retinol dehydrogenase (RDH) transforms retinol into retinal. Then RA is produced by aldehyde dehydrogenase (ALDH), a superfamily of enzymes with essential roles in detoxification and metabolism (Jackson et al. 2011). RA influences the expression of downstream target genes by binding to the RA receptors, RAR and RXR (Fig. 8A (Cunningham and Duester 2015)). Finally, RA is metabolized by the cytochrome P450 family 26 (Cyp26) enzyme, which absence of expression can restrict RA-induced responses to specific tissues or cell types (Ross and Zolfaghari 2011). Components of this pathway have been found in non-chordate animals, suggesting a more ancient origin (Canestro et al. 2006). This pathway has previously been shown to be required for *B. leachii* WBR and *Ciona* development, yet several genes required for RA signalling appear to be missing in *O. dioica* (Martí-Solans et al. 2016).

**Figure 8.**
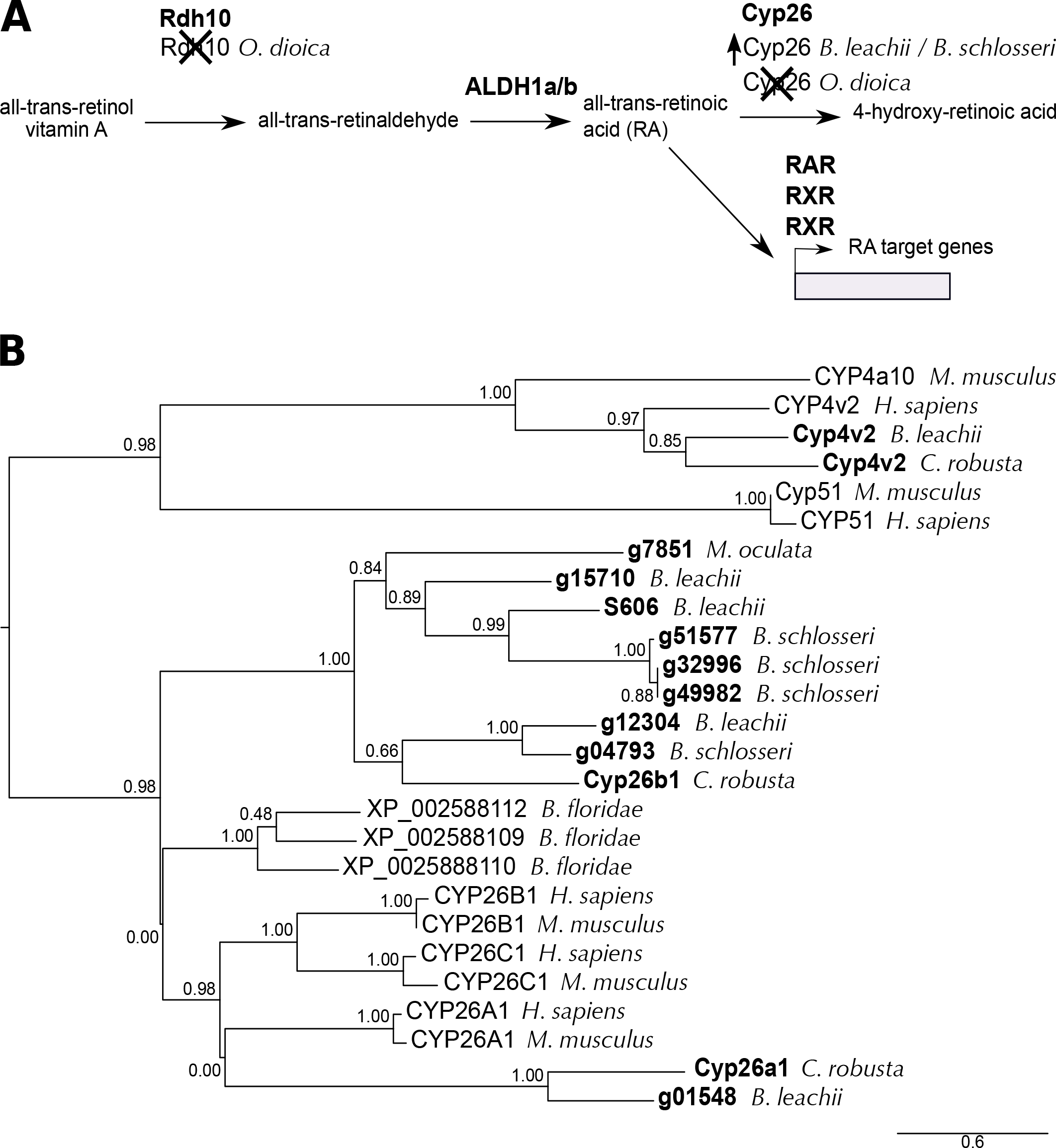
Evolution of the RA pathway in tunicates. (A) Overview of the RA synthesis and degradation pathway. In bold are the major proteins that contribute to RA signalling during animal development. Indicated below these are changes to the number of copies present in examined genomes. (B) ML phylogenetic tree depicting the relationship between invertebrate and vertebrate CYP26 proteins using CYP4 and CYP51 proteins as an out-group. Tunicate proteins are shown in bold. No *Cyp26* gene has been identified in the *O. dioica* genome. Values for the approximate likelihood-ratio test (aLRT) are indicated.

Rdh10 is the major dehydrogenase associated with the first steps of RA production, although the Rdh16 and RdhE2 enzymes can also substitute this function (Belyaeva et al. 2009; Lee et al. 2009; Belyaeva et al. 2015). The *O. dioica* genome has no orthologs for either *Rdh10* or *Rdh16* but it does have four genes that encode for RdhE2 proteins (Martí-Solans et al. 2016). *O. dioica* also lacks both an *Aldh1-*type gene as well as a *Cyp26* gene but has a single RXR-ortholog (Table S4, (Martí-Solans et al. 2016)). In contrast, the *C. robusta* genome, contains single copies of *Rdh10*, *Rdh16* and *RdhE2* genes and a total of four *Aldh1* genes, located on two chromosomes (Canestro et al. 2006). Consistent with *C. robusta*, *M. oculata*, *B. leachii* and *B. schlosseri* genomes all have single copies of *Rdh10, Rdh16* and *RdhE2* genes, as well as three *Aldh1a/b* genes on separate scaffolds (Table S4).

Three retinoic acid receptor genes were identified within the *B. leachii* genome, one of which had been cloned previously (*g03013*, (Rinkevich et al. 2007b) All three were also found in *C. robusta*, *M. oculata* and *B. schlosseri* genomes (Table S4). While there is only one potential *Cyp26* gene in *M. oculata*, four paralogs were identified in *B. leachii* and *B. schlosseri*. A phylogenetic analysis showed that these 4 genes group with CYP26 proteins (Fig. 8B, Table S4). Altogether, these results show a loss of key RA-pathway genes in *O. dioica* (*Rdh10, Rdh16, Cyp26* and *Aldh1a*), while copy numbers in other tunicate genomes increase.

## Discussion

### Genomic diversity within the Stolidobranchia

The *B. leachii* genome, along with previous genomic analyses of other ascidian species, support the widely held view that ascidian genomes are diverse and rapidly evolving, which is particularly evident in the Stolidobranchia group (Seo et al. 2001; Dehal et al. 2002b; Voskoboynik et al. 2013a; Stolfi et al. 2014; Tsagkogeorga et al. 2010; Bock et al. 2012; Tsagkogeorga et al. 2012; Rubinstein et al. 2013; Griggio et al. 2014). Nevertheless, botryllids are sufficiently similar in external appearance and morphology for early researchers to have suggested that *Botrylloides* could be a subgenus of *Botryllus* (Saito et al. 2001; Nydam et al. 2017). Strikingly however, the *B. schlosseri* genome differs from that of *B. leachii*, as well as from other sequenced tunicate genomes (Table 2). Particularly striking is the comparison between the *B. leachii* and *B. schlosseri*, where differences in genome sizes (194 Mb vs 725 Mb), the fraction of repetitive sequences (18 % vs 60 %; 65 % in (Voskoboynik et al. 2013a)) and the predicted gene number (15,839 vs 27,463; (Voskoboynik et al. 2013a)) suggest divergent genome architectures. Altogether, these comparisons indicate that the *B. schlosseri* genome has undergone a significant increase in its genomic content, including retrotransposon expansion (Table S1). In particular, there are at least two additional families in the *B. schlosseri* hAT transposon superfamily and counts of common hAT elements, such as hAT-Charlie, can differ dramatically (e.g. hAT-Charlie 366 in *B. leachii* vs 46,661 in *B. schlosseri*). DNA methylation is a key suppressor of transposon activity, changes to the methylation of transposable elements is a known driver of increased transposition (O’Neill et al. 1998; Maumus and Quesneville 2014; Simmen et al. 1999; Suzuki et al. 2007). DNA methylation in tunicate species has only been studied in *C. robusta*, and is described as mosaic, gene body methylation, whereas non-coding regions including transposons remain unmethylated (Suzuki et al. 2007), it is unknown how retrotransposons are suppressed in tunicate genomes. Nevertheless, the observed increase in transpositions could be a consequence of low non-coding DNA methylation, which may contribute to the rapid genome evolution observed in tunicate species, even between closely related species such as *B. schlosseri* and *B. leachii*.

Rapid genome evolution, and active transposable elements in particular, are proposed to aid adaptation to new environments for invasive species (Stapley et al. 2015). Differences in the colonization ability of tunicates has been noted, not only between related species such as *B. leachii* and *B. schlosseri* (Brunetti 1976; Brunetti et al. 1980; Brunetti 1974), but even at the molecular level within *B. schlosseri* populations (Bock et al. 2012; Nydam et al. 2017). It is thus possible that the observed success in tunicate invasion (Zhan et al. 2015) is supported by their plasticity in genome characteristics like transposon diversity and gene number.

Ancient homeobox gene clusters whose structure has been retained over millions of years of evolution in many organisms are fragmented in tunicate genomes. Because, the expression of each *Hox* gene across the anterior-posterior axis relates to their location within the *Hox* gene cluster (Pascual-Anaya et al. 2013), cluster breaks are predicted to have consequences for patterning processes. However, an adult body plan with correct spatial orientation of its body axes during tissue development in ascidians also needs to be established during sexual, asexual and WBR. Early patterning events in tunicate species have only been characterized during sexual reproduction in *Ciona*. Early stages of development (prior to gastrulation) follow a mosaic pattern of developmental axis formation, where inheritance of maternally provided factors establishes the body axes (Nishida 2005). *Hox* gene knockdown experiments in *C. robusta* revealed that they have very limited roles, with defects only observed in larval neuronal and tail development upon loss of *Ci-Hox11* and *Ci-Hox12* function (Ikuta et al. 2010). It thus appears that patterning events in *C. robusta* are less dependent upon anterior-posterior spatial expression of *Hox* genes to establish regional identity. Previously in *B. schlosseri*, the entry point of the connective test vessel into the developing bud determines the posterior end of the new zooid (Sabbadin et al. 1975). Therefore it is possible that ascidians incorporate environmental and physical cues to compensate for the lost gene cluster during polarity establishment. A wider analysis comprising multiple tunicate species will be necessary to investigate the exact consequences of homeobox cluster dispersion and whether the compensatory mechanism observed in *C. robusta* is the norm or an exception.

### Gene orthology analysis and coloniality candidate pathways

Among the tunicate orthologous clusters that we obtained, we identified several groups of genes that are not shared by all the tunicate genomes (Fig. 2A). Given the rapid genomic evolution of these organisms, it is more likely that these genes have either been lost or that their sequence has highly diverged, rather than independent gains of novel genes.

Of particular interest are the genes found only in the *B. schlosseri* and *B. leachii* genomes, as these may function in biological processes unique to colonial tunicates. Many of these genes have orthologs not only in vertebrates, but also in more evolutionarily distant animals such as *C. elegans* (File S4). This suggests that these genes have a more ancient origin, which was retained specifically in Botryllidae genomes. The overrepresented genes (File S4) have annotated functions including circulation (GO:0003018, GO:0003013, GO:0050880), wound healing (GO:0072378) and cell communication (GO:0007154); as well as regulation of immune cell differentiation (GO:0033081, GO:0033089), immune system process (GO:0002376) and interferons (GO:0032608). Unlike solitary tunicate species, colonial ascidians possess a complex system of single cell-lined vessels used to transport haemocytes and facilitate communication between zooids within the colony (Mukai et al. 1978). In addition, immune response is known to have roles roles in wound healing, vasculogenesis, allorecognition and regeneration (Voskoboynik et al. 2013b; Taketa et al. 2015; Gutierrez and Brown 2017; Sattler 2017). Therefore, it is possible that these genes, found only in *Botryllus* and *Botrylloides*, contribute to biological pathways and cellular processes that have important roles in colonialism.

Both *O. dioica* and *B. schlosseri* had a high number (2160 and 2716 respectively) of clusters unique to their genomes (Fig. 2A). While the *O. dioica* genome has undergone considerable loss of ancestral genes (Albalat and Cañestro 2016; Seo et al. 2001), the total number of genes in this specie is similar to that of other tunicates (Table 2). Taken together, these observations suggest that there has been a duplication of the retained genes such as *Otx* (3 copies in *O. dioica*, one in *Ciona*(Cañestro et al. 2005)), potentially involving roles in their peculiar neotenic and dioecious life cycle. The *B. schlosseri* genome has an ~10,000 higher predicted gene number compared to other tunicates (Table 2). Such massive increase in numbers suggests partial genome duplication. Further analysis will be required to determine whether these are novel or duplicated genes, hence providing important insights in the evolution of Tunicata genomes.

### Lineage-specific changes to evolutionarily conserved cell communication pathways

Cell signalling pathways are critical for morphogenesis, development and adult physiology. In particular, we have focused our analysis on three highly conserved pathways: Wnt, Notch and Retinoic Acid signalling. Representatives of all twelve *Wnt* gene subfamilies are found in metazoans, suggesting that they evolved before evolution of the bilaterians (Janssen et al. 2010). We identified members of each Wnt subfamily in tunicate genomes, along with numerous examples of lineage-specific gene loss and/or duplication. The most striking of these events was an increase in *Wnt5a* gene copy number in *B. leachii*, *B. schlosseri* and *M. oculata* genomes. Indeed, most invertebrates genomes, including the basal chordate *B. floridae*, contain a single *Wnt5* gene while most vertebrate genomes have two *Wnt5a* paralogs, believed to be a result of whole genome duplication (Martin et al. 2012). However, in the analysed tunicate genomes, up to 15 copies of this gene were identified, potentially these additional genes may have been co-opted into novel roles and were retained during tunicate evolution. Wnt5a ligands have numerous biological roles, including a suppressive one during zebrafish regeneration (Stoick-Cooper et al. 2007) and a promotive one during amphioxus regeneration (Somorjai et al. 2012). Furthermore, components of both Wnt signalling pathways are differentially expressed during WBR (Zondag et al. 2016). It is thus conceivable that *Wnt5a* gene number has expanded in colonial tunicates to sustain WBR. A functional characterization of the role of these numerous copies of Wnt5a would thus be highly interesting and potentially reveal evolutionary insights into chordate regeneration.

All components of the Notch pathway are present in the genomes we investigated. Of particular interest, the DSL Notch ligand appears to be rapidly evolving in the tunicates. This indicates that tunicate DSL proteins are under less pressure, than vertebrate orthologous proteins, to conserve their sequences. Given that the interaction between the DSL domain and the Notch receptor is core to signaling pathway activation (Chillakuri et al. 2012), it will be interesting to assess whether the functional ligand-receptor interactions between tunicate DSL proteins and tunicate Notch proteins have adapted accordingly.

Components of the RA signalling pathway have also been identified in all the tunicate genomes. However, *Oikopleura* has seemingly lost a functional RA synthesis pathway, while still forming a functional body plan. This suggests that either uniquely RA is not involved in critical developmental events in this species, that the RA signalling function has been replaced or that *O. dioica* utilizes an alternative synthesis approach. Conversely, lineage specific increases in RA pathway gene numbers have been observed in *C. robusta* (Aldh1, (Sobreira et al. 2011)) and *Botrylloides* (*CYP26* genes, Fig. 8).

RA, Notch and Wnt pathways play roles in regeneration and development in many species, including Stolidobranchian tunicates (Rinkevich et al. 2007b, 2008; Zondag et al. 2016) and *Cionidae* (Hamada et al. 2015; Jeffery 2015). The observed loss of RA signalling genes may result in reduced regeneration ability for *O. dioica*, however it’s regenerative abilities have not been characterized. Given the unique chordate WBR potential developed by colonial tunicates, it is conceivable that there is selective pressure on their genomes to retain these pathways. We thus predict that these pathways play a similar role in colony reactivation following hibernation.

Among tunicates there exist significant differences in life cycle, reproduction and regeneration ability, even between closely related species of the same family, which likely reflect an underlying diversity in genomic content. For instance, differences in both asexual and sexual reproduction have been observed within the Botryllidae family (Oka and Watanabe 1957; Brunetti 1974, 1976, Berrill 1951, 1947, 1941). Furthermore, *B. schlosseri* can only undergo WBR during a short time frame of their asexual reproductive cycle when the adults are reabsorbed by the colony (Voskoboynik et al. 2007; Kürn et al. 2011) while *B. leachii* can undergo WBR throughout their adult life (Rinkevich et al. 2007b). Overall, this indicates that despite a generally similar appearance, the rapid evolution of the Tunicata subphylum has provided diversity and innovations within its species. It will be interesting to investigate how such genomic plasticity balances between adaptation to new challenges and constraint, preserving common morphological features, in future studies.

In conclusion, our assembly of the *B. leachii* genome provides an essential resource for the study of this colonial ascidian as well as a crucial point of comparison to gain further insights into the remarkable genetic diversity among tunicate species. In addition, the genome of *B. leachii* will be most useful for dissecting WBR in chordates; particularly through comparison with *B. schlosseri* for understanding how the initiation of WBR can be blocked during specific periods of their life cycle. Furthermore, given the key phylogenetic position of Tunicates with respect to vertebrates, the analysis of their genomes will provide important insights in the emergence of chordate traits and the origin of vertebrates.

## Methods

### Sampling, library preparation and sequencing

*B. leachii* colonies were collected from Nelson harbour (latitude 41.26°S, longitude 173.28°E) in New Zealand. To reduce the likelihood of contamination, embryos were dissected out of a colony and pooled before carrying out DNA extraction using E.Z.N.A SP Plant DNA Mini Kit. A total of 2 µg each was sent to New Zealand Genomics Limited (NZGL) for library preparation and sequencing. Short read sequencing of Illumina TruSeq libraries in a HiSeq2500 generated 19,090,212 paired-end reads of 100 bp (average fragment size: 450 bp, adaptor length: 120 bp). A second sequencing (Illumina Nextera MiSeq Mate Pair) not size-selected generated 31,780,788 paired-end sequences of 250 bp (fragment size: 1.5 - 15 kb, median size: ~3 kb, adaptor length: 38 bp).

PreQC report was generated using the String Graph Assembler software package (Simpson 2014) and quality metrics before assembly with both FastQC (Andrews 2010) as well as MultiQC (Ewels et al. 2016) (Fig. S1). These analyses revealed that 91 % of sequences had a mean Phred quality score >= 30, 96 % of bases a mean Phred quality score >= 30, and 39 % of sequences an adapter sequence (either Illumina or Nextera). Adaptor trimming was performed with NxTrim (O’Connell et al. 2015) for the mate pair library, followed by Trimmomatic (Bolger et al. 2014) with the following options: MINLEN:40 ILLUMINACLIP:2:30:12:1:true LEADING:3 TRAILING:3 MAXINFO:40:0.4 MINLEN:40 for both libraries. After trimming, 86,644,308 paired-end (85 %) and 12,112,004 (12 %) single-end sequences remained (100 % with a mean Phred quality score >= 30, < 1 % with an adapter sequence).

### Genome assembly

*De novo* assembly was performed in three consecutive iterations following a Meta-assembly approach (Table S5). First, both libraries were assembled together in parallel, using a k-mer size of 63 following the results from KmerGenie (Chikhi and Medvedev 2014) whenever available, by five assemblers: AbySS (Simpson et al. 2009), Velvet (Zerbino and Birney 2008), SOAPdenovo2 (Luo et al. 2012), ALLPATHS-LG (Gnerre et al. 2011), MaSuRCA (Zimin et al. 2013). The MaSuRCA assembler was run twice, once running the adapter filtering function (here termed “MaSuRCA-filtered”), the other without (termed simply “MaSuRCA”). Their respective quality was then estimated using three different metrics: the N50 length, the BUSCO core-genes completion (Simão et al. 2015) and the Glimmer number of predicted genes (Delcher et al. 1999). Second, these drafts were combined by following each ranking using Metassembler (Wences and Schatz 2015), hence producing three new assemblies (limiting the maximum insert size at 15 kb). Third, the *B. leachii* transcriptome (Zondag et al. 2016) was aligned to each meta-assembly using STAR (Dobin et al. 2013), which were then combined thrice more using Metassembler following their alignment percentage and limiting the maximum insert size at 3 kb, 8 kb and 15 kb. Finally, the quality of the meta-meta-assemblies was estimated using the BUSCO score and the best one (Table S5) selected as the reference *de novo* assembly.

### Data access

All data was retrieved from the indicated sources in January 2016. Note that *Ciona intestinalis* type A (Dehal et al. 2002b) has recently been recognized as a distinct species (*Ciona robusta*, (Brunetti et al. 2015)) and that this study has been undertaken before it was renamed.

*B. leachii*, *B. schlosseri*, *C. robusta*, *M. oculata*: Ascidian Network for *In Situ* Expression and Embryonic Data (ANISEED, https://www.aniseed.cnrs.fr/aniseed/, (Tassy et al. 2010)). *O. dioica*: Oikopleura Genome Browser (http://www.genoscope.cns.fr/externe/GenomeBrowser/Oikopleura/, (Seo et al. 2001)). *B. floridae*, *H. sapiens*: Joint Genome Institute (JGI, http://genome.jgi.doe.gov, (Grigoriev et al. 2012))

#### Repeat region analysis

A *de novo* repeat library was build for each tunicate genome using RepeatModeler (Smit and Hubley 2015). This utilizes the RECON tandem repeats finder from the RepeatScout packages to identify species-specific repeats in a genome assembly. RepeatMasker (Smit et al. 2015) was then used to mask those repeats.

### Gene annotation

*Ab initio* genome annotation was performed using MAKER2 (Holt and Yandell 2011) with Augustus (Stanke and Waack 2003) and SNAP (Korf 2004) for gene prediction. In addition, we used our previously published transcriptome (Zondag et al. 2016) and a concatenation of UniProtKB (UniProt Consortium 2015), *C. robusta* and *B. schlosseri* proteins into a custom proteome as evidence of gene product. Using the predicted genes, Augustus and SNAP were then trained to the specificity of *B. leachii* genome. A second round of predictions was then performed, followed by a second round of training. The final annotation of the genome was obtained after running a third round of predictions, and the provided trained Augustus and SNAP configurations after a third round of training. Non-coding RNA sequences were then annotated using Infernal (Nawrocki and Eddy 2013) with Rfam library 12.0 (Nawrocki et al. 2015), tRNAscan-SE (Lowe and Eddy 1997) and snoRNA (Lowe 1999). Finally, the identified sequences were characterized by InterProScan (Jones et al. 2014).

### Analysis of Gene Ontology terms

Distribution of Gene Ontology (GO) terms were computed for each species as follows. GO terms were extracted from the genome annotation and the number of occurrence for each term determined using a custom Python script. The resulting list of frequencies was then simplified using REVIGO (similarity factor “small” of 0.5, (Supek et al. 2011)) and the TreeMap output retrieved. The hierarchy of every GO term present was reconstructed following the schema defined by the core gene ontology (go.obo, (The Gene Ontology Consortium 2015)) using a custom Python script selecting the shortest path to the root of the tree, favouring smaller GO terms identification number in case of multiple paths. Finally, frequencies were displayed using the sunburstR function of the Data-Driven Documents library (D3, (Bostock et al. 2011)).

Predicted amino-acid sequences for all species were retrieved and clustered into 17,710 groups by OrthoMCL (Li et al. 2003). Protein sequences within each group were then aligned into a Multiple Sequence Alignment (MSA) by Clustal-Omega, and the corresponding consensus sequence inferred by EMBOSS cons. Consensus sequences were matched to the Swiss-Prot curated database using BLASTp (e-value cut-off of 10^-5^), and the GO terms corresponding to the best match retrieved. GO terms frequencies were analysed as described above and displayed using REVIGO’s treemap. The overrepresentation analysis was performed using GOrilla (Eden et al. 2009) with *Homo sapiens* as the organism background, using a *p*-value threshold of 10^-3^ and REVIGO treemap (similarity factor “medium” of 0.7) for visualization.

### Analysis of specific gene families

Genes and transcripts for each examined genome were identified by a tBLASTn search with an e-value cut-off at 10^-5^ using the SequencerServer software (Priyam et al. 2015). This was followed by a reciprocal BLAST using SmartBLAST (NCBI Resource Coordinators 2016), to confirm their identity.

Delta serrate ligand conserved protein domain (PF01414) was used to identify the corresponding proteins in tunicate genomes. To identify *Notch* receptor genes the conserved LNR (lin-notch repeat) domain (PF00066) was used. ALDH-like genes were identified by tBLASTn search (PF00171) and classified using SMART blast.

### Phylogenetics

Sequences were aligned with ClustalX (Jeanmougin et al. 1998) before using ProtTest 3 (Abascal et al. 2005) to determine the best-fit model of evolution. The best-fit model for the DSL phylogeny was WAG+I+G and, for CYP26 proteins, was LG+I+G.

Bayesian inference (BI) phylogenies were constructed using MrBayes (Ronquist and Huelsenbeck 2003) with a mixed model for 100,000 generations and summarized using a Sump burn-in of 200. Maximum Likelihood (ML) phylogenies were generated by PhyML (Guindon et al. 2010), using the estimated amino acid frequencies.

Accession numbers are provided in File S3 and sequence alignments are provided in Figure S3. Analyses carried out with BI and ML produced identical tree topologies. Trees were displayed using FigTree v1.4.2 (Rambaut 2016).

## Acknowledgements

Funding support was provided to M.J.W. by the Otago BMS Deans Bequest and Department of Anatomy. S.B. was supported by the Swiss National Science Foundation (SNSF) grant number P2ELP3_158873. We would like to thank Peter Maxwell and the New Zealand eScience Infrastructure (NeSI); Christelle Dantec and ANISEED for help and advice during the annotation process, as well as for the accompanying *B. leachii* genome browser.

## Supplementary Figures

**Fig. S1**. SGA analysis. Including genome size estimation and de Brujin graph quantification.

**Fig. S2**. Gene Ontology terms identified in the larger orthologs clusters between the tunicate genomes.

**Fig. S3**. Protein sequence alignments used to generate the Notch and RA phylogenies.

### Supplementary Files

**File S1.** BUSCO scores for the *B. leachii* genome assembly

**File S2.** Results of Repeatmasker analysis using de novo repeat libraries (Repeatmodeler)

**File S3.** Results from OrthoMCL group including the REVIGO GO term listed each orthologue group, used in the comparison of the GO terms between the tunicate genomes.

**File S4.** BLASTp, GOrillia and REVIGO results used used for overrepresentation analysis of GO terms present in the *B. leachii* and *B. schlosseri* only orthologue group analysis.

**File S5.** Corresponding Gene and Transcript IDs for *B. leachii* genes of interest. Accession numbers for protein sequences used in phylogeny construction.

### Supplementary Tables

**Table S1**. Repetitive elements identified in the *B. leachii* and *B. schlosseri* genomes using Repeatmoduler and RepeatMasker.

**Table S2**. Comparison of the number of Wnt pathway genes.

**Table S3**. Comparison of the number of Notch pathway genes.

**Table S4**. RA pathway components across annotated tunicate genomes.

**Table S5**. Iterative results of the meta-assembly approach followed for the *de novo* assembly of the *B. leachii* genome

